# Cranial anatomy of the predatory actinopterygian *Brazilichthys macrognathus* from the Permian (Cisuralian) Pedra de Fogo Formation, Parnaíba Basin, Brazil

**DOI:** 10.1101/540310

**Authors:** Rodrigo T. Figueroa, Matt Friedman, Valéria Gallo

## Abstract

*Brazilichthys macrognathus* is the only named actinopterygian from the Permain (Cisuralian) Pedra de Fogo Formation of northeastern Brazil, where it is represented by a single three-dimensionally preserved but incompletely described skull of unclear systematic placement. We used X-ray computed microtomography (μ-CT) to better document its anatomy and phylogenetic affinities. μ-CT reveals parts of the internal skeleton. We correct errors in original description, including the number of infraorbital bones and the misidentification of the dermosphenotic as sclerotic ossifications. These reinterpretations of external anatomy are joined by new data on internal structure, including the palate, parasphenoid, and branchial and hyoid arches. A maximum parsimony analysis of anatomical data resolves *Brazilichthys* as a stem actinopterygian, crownward of all Devonian species. This placement is supported by the absence of a dermosphenotic posterior ramus and the presence of opercular process of the hyomandibula. A similar placement is suggested by a Bayesian analysis of this same dataset, although relationships throughout the tree are less resolved. Our results reject previous interpretations of *Brazilichthys* as a relative of Birgeriidae, a Triassic group consistently placed within the actinopterygian crown. Although *Acrolepis* is too poorly known to be included in our analysis, we also reject a close relationship between this taxon and *Brazilichthys*, as their only shared similarities appear to be broadly distributed among early actinopterygians.

## INTRODUCTION

The late Paleozoic is an important episode of actinopterygian evolution, representing a bridge from low-disparity and low-diversity ray-finned fish faunas of the Devonian to the emergence of the earliest teleosts in the Triassic (Sallan, 2014; Friedman, 2015). This Permo-Carboniferous interval is marked by substantial innovation in skull and body morphology (Sallan and Friedman, 2012), as well as the possible origin of the crown radiation and divergence of the cladistian, chondrostean, and neopterygian total groups (Giles et al., 2017). Despite its clear significance, the Permo-Carboniferous remains a poorly known interval in the fossil record of fishes. Despite relatively abundant actinopterygian fossils during some parts of the late Paleozoic, their taxonomy is confused and relatively few species are known in detail. Consequently, fossil fishes of Carboniferous and Permian age are among the least stable taxa in analyses of actinopterygian interrelationships (Giles et al., 2017). Compounding these issues, the known Permo-Carboniferous record shows a strong geographic collecting and research bias toward northern landmasses, with the best known actinopterygian faunas of this age deriving from North American (Mississippian: Bear Gulch; Pennsylvanian: Mazon Creek, Linton, Kinney Brick Quarry; Schultze and Bardack, 1987; Kues and Lucas, 1992; Hansen, 1996; Poplin and Lund, 2002) and European (Mississippian: Foulden, Wardie, Bearsden; Pennsylvanian: Bohemian Massif; Permian: East Greenland, Zechstein; Aldinger, 1937; Gardiner, 1985; Haubold and Schaumberg, 1985; Coates, 1998; Štamberg, 2013) localities. While a handful of productive localities are known from southern continents, these have generally been the subject of broad faunal overviews (e.g. Witteburg Group of South Africa; Gardiner, 1969) or detailed descriptions of only single constituent taxa (e.g. *Ebenaqua* from Rangal Coal Measures of Blackwater, Australia; Campbell and Phuoc, 1983).

Brazilian deposits yield the vast majority of Permo-Carboniferous actinopterygians known from South America (Cione et al., 2010), with only a handful of examples known from elsewhere, mostly based on poorly preserved specimens (e.g. Beltan, 1978; this material is now considered lost, pers. comm. Piñeiro, G., April 18, 2017). Despite the relative neglect of the South American record of Paleozoic fishes, sporadic research efforts reveal substantial assortment of Permian actinopterygians from Brazil. These span the Permian and overwhelmingly derive from deposits in the Paraná Basin of southern Brazil: the Rio do Sul (Cisuralian in age and yielding *Elonichthys gondwanus*; Richter et al., 1985), Campo Mourão (Cisuralian in age and yielding *Roslerichthys riomafrensis* and *Santosichthys mafrensis*; Hamel, 2005; Malabarba, 1988), Rio Bonito Formation (Guadalupian-Lopingian in age yielding *Tholonosteon santacatarinae*; Richter et al., 1985) Rio do Rasto (Guadalupian-Lopingian in age and yielding *Rubidus pascoalensis* and *Paranaichthys longianalis*; Richter, 2002; Dias, 2012), and Corumbataí (Lopingian in age and yeilding *Tholonotus brasiliensis* and *Angatubichthys mendesi*; Dunkle and Schaeffer, 1956; Figueiredo and Carvalho, 2004) formations. By contrast, *Brazilichthys macrognathus* is the only Permian actinopterygian known from the Parnaíba Basin in northeastern Brazil (Figueroa and Machado, 2018). *Brazilichthys* in many ways encapsulates the problems surrounding the study of Permo-Carboniferous fishes from Brazil and elsewhere. Known only from the holotype specimen, *Brazilichthys* has only been described externally (Cox and Hutchinson, 1991). The limited data available for the genus have led to informal alignment with multiple lineages of early actinopterygians on the basis of overall resemblance: first Acrolepidae by Cox and Hutchinson (1991), and then Birgeriidae by Romano and Brinkmann (2009).

Some of the ambiguity surrounding the phylogenetic placement of many early ray-finned fishes stems from restricted anatomical description, often restricted to superficial details of the dermal skeleton. The widespread availability of micro computed tomography (μ-CT) now permits detailed examination of character-rich internal skeletal features. Application of μ-CT to previously described early actinopterygians has resulted in substantial new information for previously described taxa that has helped to refine--and in some cases substantially change–their inferred phylogenetic positions (Giles et al., 2017; Argyriou et al., 2018; Coates and Tietjen, 2019; Friedman et al., in 2019). Here we provide a revised description of *Brazilichthys macrognathus* based on μ-CT scans of the type and only specimen. Using these new data in combination with a recently developed character matrix, we examine the phylogenetic placement of *Brazilichthys* among early actinopterygians, comparing our results to previous hypothesis regarding the actinopterygian stem. Further, we compare *Brazilichthys* to large predatory Paleozoic actinopterygians, proposing steps for future studies.

## MATERIALS AND METHODS

### Specimens Examined

*Brazilichthys macrognathus*, holotype, DGM 1061-P (Fig. 1A-C).

**FIGURE 1.**
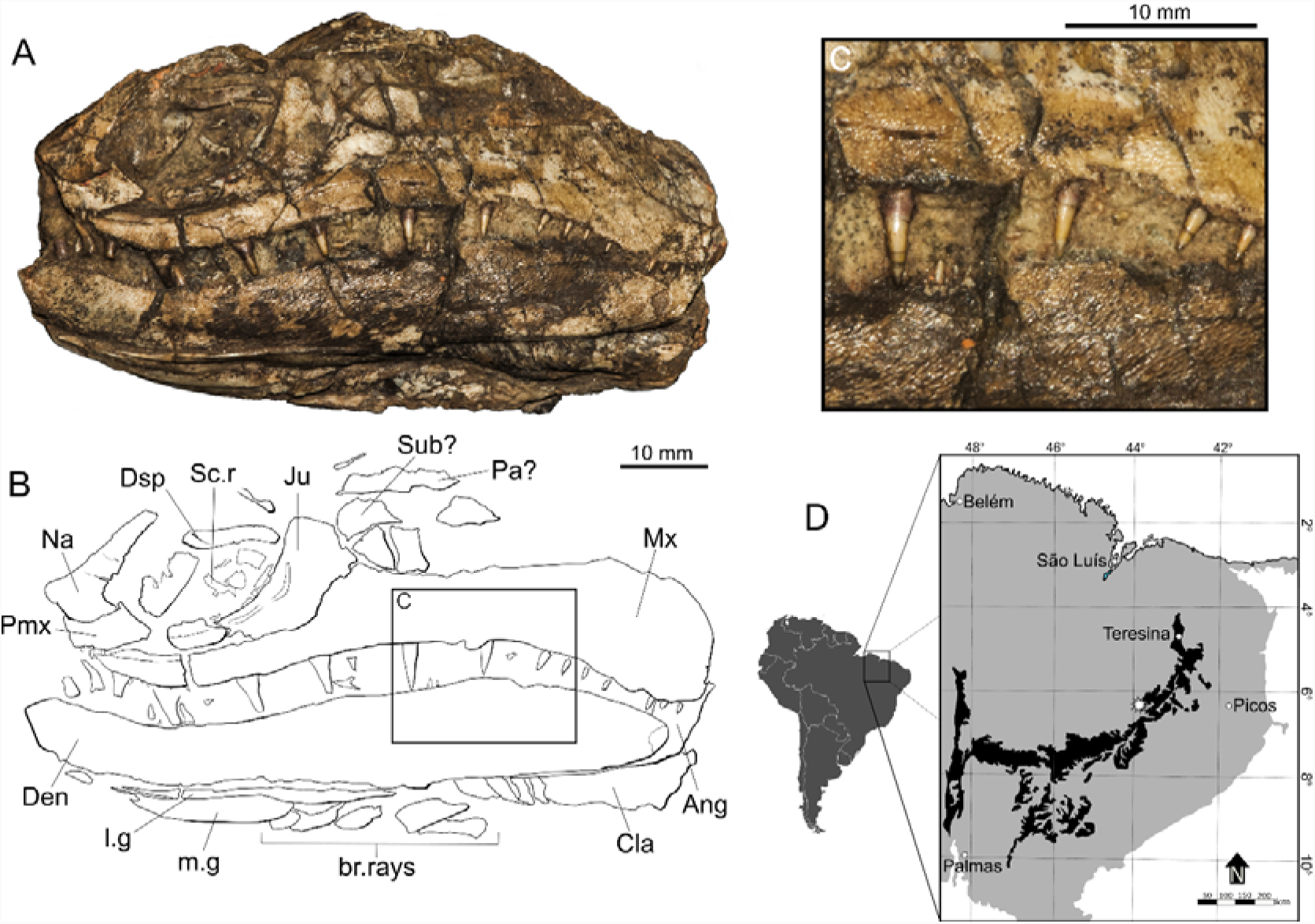
*Brazilichthys macrognathus*, DGM 1061-P, holotype, Pedra de Fogo Formation, Pastos Bons locality, state of Maranhão, Brazil. **A**, Specimen photograph in left lateral view. **B**, Interpretive drawing. **C**, Close-up of maxilla showing fine ornament ridges and acrodin caps on teeth; **D**, Type locality within the Pedra de Fogo Formation (black) of the Parnaíba Basin (grey), based on Santos and Carvalho (2009).

### Computed Microtomography

Computed microtomography (μ-CT) of DGM 1061-P was conducted at the Laboratório de Instrumentação Nuclear, of the Instituto Alberto Luiz Coimbra de Pós-graduação e Pesquisa de Engenharia (COPPE), located in the Centro de Tecnologia (CT) of the Universidade Federal do Rio de Janeiro (UFRJ), using a SkyScan 1173 scanner.

Parameters of the scan were: current = 61 μA; voltage =130 kV; projections = 2234. A 1 mm copper filter was used and projections were processed in in the proprietary software NRecon 1.6.9.4 to produce a tomogram stack. The resolution of the scan was 35.61 μm.

Segmentation was completed in Spiersedit 2.20 (Smith et al., 2016), using tomograms saved as .bmp format. Downsampling (by 50% in x, y, and z axes) of the data was done to speed the construction of the 3D model, without any conspicuous loss of detail. The slices were then processed manually, with the resulting 3D model was initially visualized using Spiersview 2.20. More minor modifications such as smoothing and brightness were made using this software, along with removal of ‘islands’ of sediment or unidentifiable bone fragments. Production of final images was completed in Blender (blender.org) (Garwood and Dunlop, 2014). Illustrations of the resulting renders and specimen reconstructions were completed in Inkscape (inkscape.org; Harrington, 2005). Blender models exported as .ply files and the tomographic stack are publically available through the following link: https://doi.org/10.6084/m9.figshare.7600103.

### Phylogenetic Dataset

*Brazilichthys macrognathus* was coded for the characters in the matrix presented by Giles et al. (2017). *Brachydegma caelatum* was excluded from this matrix because its anatomy is under revision by M. Friedman and others, and available descriptions are not reliable. The complete matrix contains 93 taxa and 265 unweighted characters (Supplementary Data 1). *Brazilichthys* can be evaluated for 33% of all characters. The analysis includes several non-actinopterygian fishes, with *Dicksonosteus arcticus* set as the outgroup. In contrast to Giles et al. (2017), we did not enforce relationships among non-actinopterygian taxa. All characters were treated as unordered in both analysis.

#### Parsimony Analysis

An equally weighted parsimony analysis was conducted using the software TNT 1.5 (Goloboff et al., 2016). The New Technology search algorithm was used, with 5 initial additional sequences. Bremer support was calculated in TNT using the TBR (“tree bisection and reconnection”) for all most parsimonious trees. The results where then plotted against the strict consensus tree. Nodes with Bremer support below 1 were automatically collapsed. A formatted file is provided in Supplementary Data 2 to reproduce results of the parsimony analysis.

The most parsimonious trees (Supplementary Data 3) were used to calculate the strict consensus topology, and the file was exported to Mesquite (Maddison and Maddison, 2018) where we mapped the evolution of each character based on the likelihood and parsimony algorithms (Lewis, 2001). The strict consensus tree with the character mapping is available through the link: https://doi.org/10.6084/m9.figshare.7600103.

#### Bayesian Analysis

The Bayesian analysis was conducted on MrBayes 3.2.5 (Ronquist et al., 2012) using the Metropolis Coupled Markov Chain Algorithm (MCMC) and the MkV model for discrete morphological data (Lewis, 2001; Wright and Hillis, 2014). Character coding was set to “variable” and a gamma distribution was incorporated. The number of substitution types was set to “nst2”, which mean that all transitions have potentially different rates. The number of generations was initially set to 500,000, with the number of generations increased until reaching a low standard deviation of split frequencies. Burn-in fraction was set to 50% of the resulting topologies. A complete script for MrBayes is given in Supplementary Data 4.

The resulting phylogram (Supplementary Data 5) was visualized in FigTree 1.4.3 (Rambaut, 2018) and the tree file was exported to Mesquite (Maddison and Maddison, 2018) where we mapped the evolution of each character based on the likelihood and parsimony algorithms (Lewis, 2001). The Bayesian consensus with all mapped synapomorphies is available through the link: https://doi.org/10.6084/m9.figshare.7600103.

### Terminological Conventions

Following McCune and Schaeffer (1986; see also Friedman and Giles 2016), we apply the term ‘paleopterygian’ to designate the assemblage of Paleozoic ray-finned fishes of uncertain relationships to one another and modern actinopterygian clades. Our use of this term is not a suggestion that species falling within this category represent a natural group, as we explicitly seek to avoid the taxonomic connotations associated with baggage-laden terms like ‘palaeonoiscoid’ or ‘palaeonisciform’. This nomenclature is plastic and with no implication of evolutionary affinities, although it is our anticipation that groups will be extracted from this paleopterygian assemblage as their relationships to fossil and living actinopterygians are clarified with further anatomical and phylogenetic investigation.

Bone nomenclature adopted here follows the conventional terminology for actinopterygians as in Gardiner (1984). We acknowledge that the frontals and parietals of actinopterygians under this scheme are the homologues of sarcopterygian pareitals and postparietals, respectively (Westoll, 1943; Schultze and Arsenault, 1985).

#### Anatomical Abbreviations

**Ac.Vo**, accessory vomer; **Ang**, angular; **app**, anterior process of parasphenoid; **Art**, articular; **asp**, ascending process of parasphenoid; **B.rays**, branchiostegal rays; **bpt**, basipterygoid process; **Cbr**, ceratobranchial; **Chy**, ceratohyal; **Cla**, clavicle; **Cor**, coronoid; **Den**, dentary; **Dhy**, dermohyal; **Dsp**, dermosphenotic; **e.g**, extralateral gular; **Epb**, epibranchial; **f**, foramen; **f.mand.V**, foramen for mandibular branch of trigeminal nerve; **f.scl,** sensory canal foramen; **fo.add**, fossa for adductor muscle; **h.op**, opercular process of hyomandibula; **Hb**, hypobranchials; **Hb_1_**, first hypobranchial; **Hh**, hypohyal; **Hy**, hyomandibula; **hy.c**, hyomandibular canal; **i.sc**, infraorbital sensory canal; **Inf**, infraorbitals; **Inf.s**, infraorbital series; **Ju**, jugal; **l.g**, lateral gular; **m.g**, median gular; **Men**, mentomeckelian; **Mx**, maxilla; **n.ao**, anterior nasal opening; **n.po**, posterior nasal opening; **Na**, nasal; **p.l**, pit-line; **Pa**, parietal; **pal.te**, palatal teeth; **Part**, prearticular; **Pmx**, premaxilla; **Pop**, preoperculum; **Pq**, palatoquadrate; **Propt**, propterygium; **Psp**, parasphenoid; **San**, surangular; **Sc.r**, sclerotic ring; **scl**, sensory canal line; **Sob**, supraorbital; **spig**, spiracular groove; **Sub**, suborbital; **Te**, teeth; **V**, foramen or canal for trigeminal nerve.

SYSTEMATIC PALEONTOLOGY

ACTINOPTERYGII Cope, 1887

BRAZILICHTHYIDAE Cox & Hutchinson, 1991

*BRAZILICHTHYS* Cox & Hutchinson, 1991

*BRAZILICHTHYS MACROGNATHUS* Cox & Hutchinson, 1991

#### Type and Only Specimen

DGM 1061-P, incomplete skull. The specimen is housed at the paleontological collection of the Museu de Ciências da Terra (MCT), of the Departamento Nacional de Produção Mineral (DNPM), on behalf of the Centro de Pesquisas de Recursos Minerais (CPRM)

#### Type Locality and Horizon

Pedra de Fogo Formation, Pastos Bons Locality (∼ 6° 40’ S, 44° 04’ W), between the cities of Pastos Bons and Nova Iorque, state of Maranhão, Brazil (Fig. 1D). The Pedra de Fogo Formation is assigned to the Artinskian-Kungurian stages of the Permian based on its palynological assemblage (Iannuzzi et al., 2018).

#### Emended Diagnosis

‘Paleopterygian’ actinopterygian distinguished by the following combination of characters: parasymphysial fangs, some of which are strongly procumbent; flexed symphysial region of the mandible in lateral view; widely spaced glenoid fossae of articular; long ellipsoidal median and lateral gulars; presence of long extralateral gulars; rod-like dermosphenotic; parasphenoid with distinct basipterygoid processes, short and rectangular ascending processes and robust anterior process, and a prominent dorsal keel on the anterior corpus.

#### Notes

The bones originally mentioned by Cox and Hutchinson (1991) were: nasal, rostral, pre-maxilla, infraorbito-suborbital, infraorbital (jugal), infraorbital (lacrimal?), suborbital, maxilla, dentary, angular, clavicle, median gular, lateral gulars, branchiostegal rays, and part of the sclerotic ring. However, the holotype of *B. macrognathus* lacks the rostral bone described by Cox & Hutchinson (1991), suggesting loss or other damage in the time between their account and this redescription. Cox & Hutchinson (1991) also illustrated the delicate ornamentation of the dermal bones of the skull, which is composed of closely spaced wavy ridges.

### Description

#### Skull Roof

The skull roof is represented only by isolated fragments visible dorsal to the circumorbital series. A small fragment dorsal to the dermosphenotic might represent part of the frontal (Figs 2,3), but it lacks any diagnostic features (e.g. sensory canals). A posterior element above the suborbital might be part of the left parietal bone, but this too lacks any characteristic detail.

**FIGURE 2.**
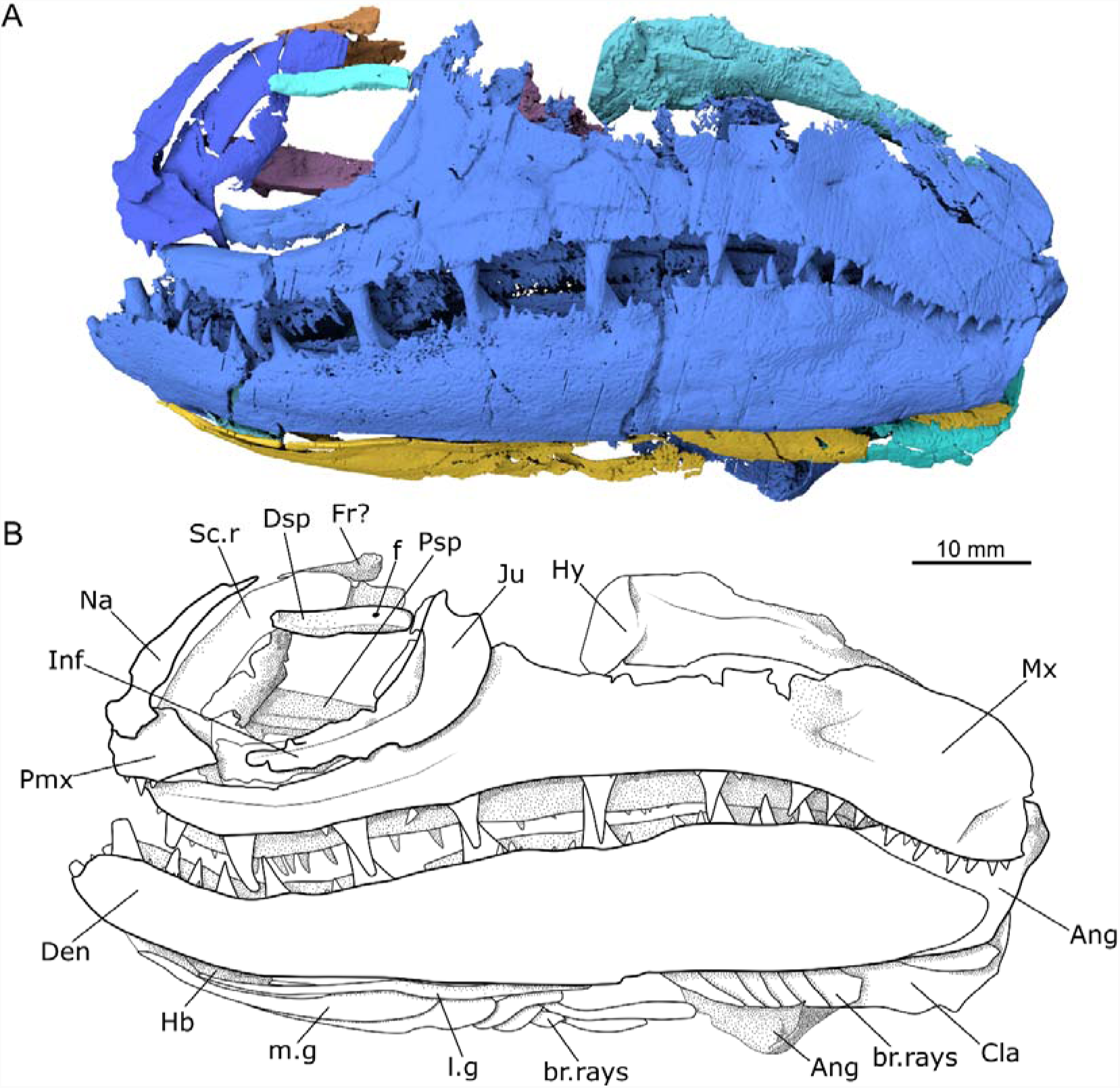
*Brazilichthys macrognathus*, DGM 1061-P, skull in left-lateral view. **A**, Surface rendering based on μ-CT data; **B**, Interpretive drawing.

**FIGURE 3.**
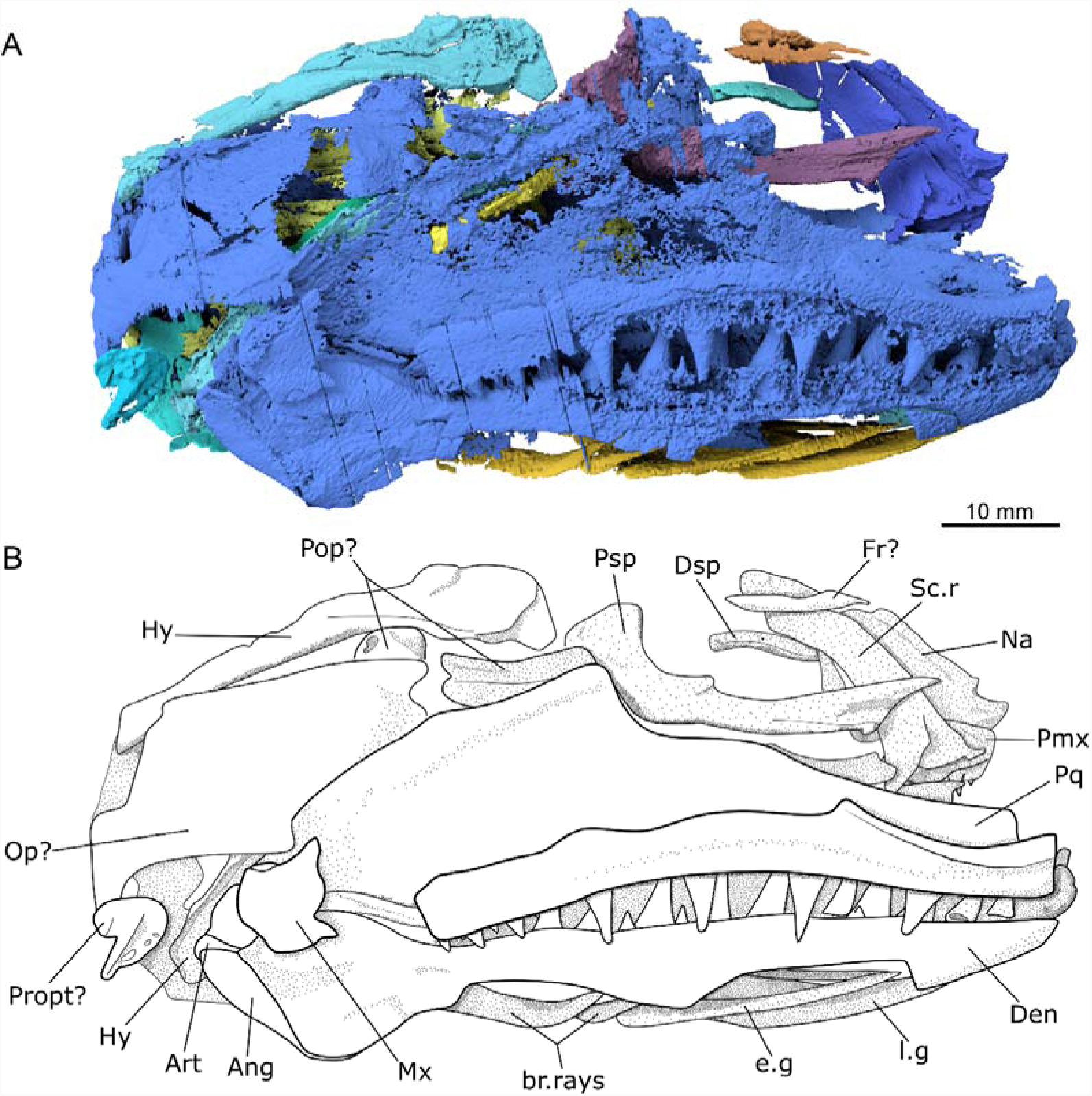
*Brazilichthys macrognathus*, DGM 1061-P, skull in right-lateral view. **A**, Surface rendering based on μ-CT data; **B**, Interpretive drawing.

#### Snout Region and Circumorbital Series

The nasal is large and bears notches for the anterior and posterior nasal openings (Fig. 4). There is no obvious indication of the sensory canal externally, but it is apparent internally as a slight longitudinal groove near the anterior border of the bone. The positioning of this groove is consistent with the sensory canal line of the nasals of early actinopterygians. The anterior nasal opening would have been enclosed anteriorly by the the rostral, which is not preserved but was reported in a previous description (Cox & Hutchinson, 1991). The posterior nasal opening is confluent with the orbital opening.

**FIGURE 4.**
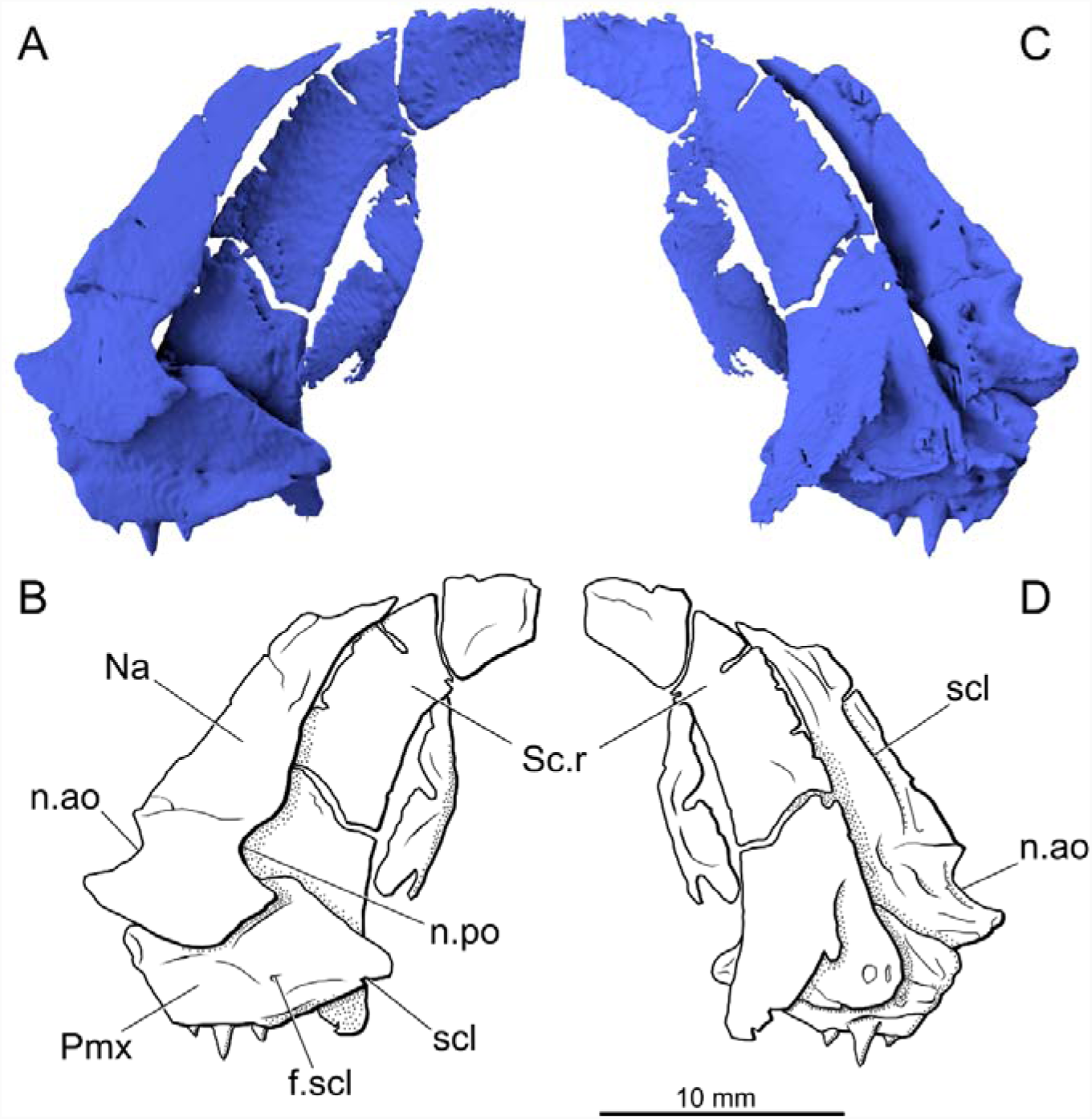
*Brazilichthys macrognathus*, DGM 1061-P, left antorbital region. **A**, Surface rendering in external view based on μ-CT data; **B**, interpretive drawing; **C**, surface rendering in internal view based on μ-CT data; **D**, interpretive drawing.

A slender bone located inside the orbital cavity of the DGM 1061-P was originally identified as part of the sclerotic ring. However, μ-CT revealed an enclosed canal extending through half of the length of the bone, exiting by a foramen in the center of the outer surface (Figs. 2,3). Bones of the sclerotic ring are not canal bearing, indicating a different affinity than that proposed by Cox & Hutchinson (1991). Due to the peculiar path drawn by this canal and its positioning, the bone is interpreted here as the dermosphenotic. It is an elongate robust bone that contacts the nasal anteriorly, the jugal posteriorly, and probably the frontal dorsally, composing the dorsal margin of the orbit.

The sclerotic ring is partially visible superficially on the specimen. However, a large, thin element mesial to the nasal and premaxilla represents a concealed part of the sclerotic ring (Fig. 4). This element is large in comparison to the orbital opening but its outline closely matches it in shape.

The rest of the circumorbital series is well preserved. The infraorbital series is composed of three infraorbitals. These are, from anterior to posterior: the lachrymal, a single infraorbital, and the jugal. The jugal is lunate and the infraorbital sensory canal line lies near its anterior border, without any evidence of posterior branching. This bone slightly overlaps the postorbital expansion of the maxilla. The infraorbital lies anterior to the jugal and bears the extension of the infraorbital sensory canal line. The lachrymal is displaced within the orbital cavity but would contact the premaxilla and the nasal anteriorly and the infraorbital posteriorly. A poorly preserved rhomboidal bone lies ventral to the jugal and the posterior expansion of the maxilla. Cox & Hutchinson (1991) interpreted it as a suborbital, an identification adopted here due to the absence of any sensory canals.

The premaxilla is a short and robust bone. Its triangular posterior border, restricted to the anterior margin of the orbit. It bears an enclosed sensory canal that exits to the surface medially by small foramina on the exposed surface of the premaxilla. It is partially overlapped posterodorsally by the nasal. We assume it would contact the (unpreserved) rostral anterodorsally. The premaxilla bears at least 3 small conical teeth.

#### Jaws and Palate

Both the dentary and the maxilla bear two series of teeth (Figs. 5), one inner series composed of large conical and posteriorly directed teeth and one outer series of much smaller teeth. Tooth rows extend the complete length of these bones, even on the portion of the maxilla that overlaps the dentary. The teeth in this area are anteriorly directed. The big teeth of the lingual series bear acrodin tooth caps (Fig. 1C).

**FIGURE 5.**
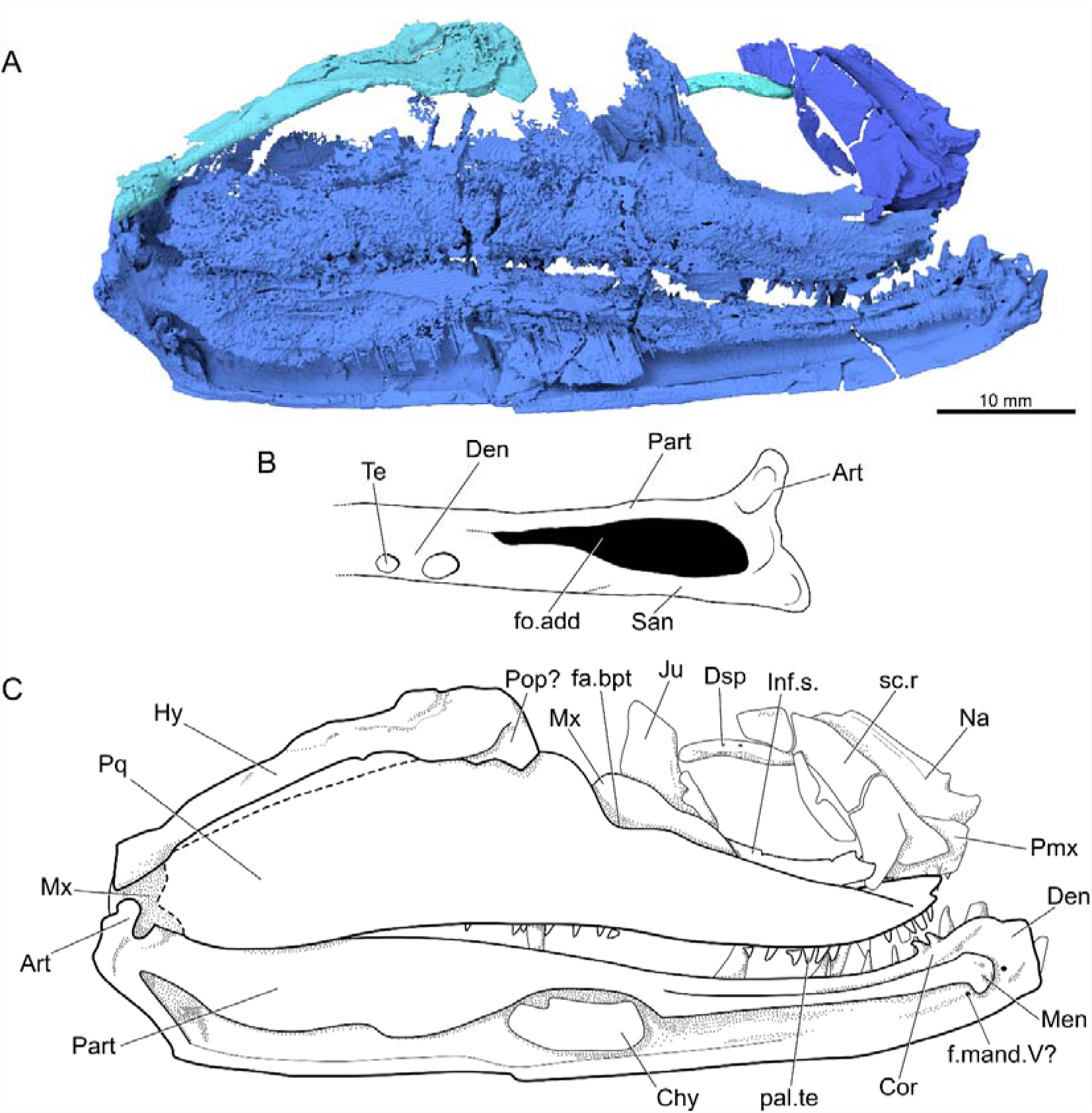
*Brazilichthys macrognathus*, DGM 1061-P, left palate, lower jaw, hyomandibula, and regions of dermal skull in mesial view. **A**, surface render based on μ-CT data; **B**, dorsal view of articular region and adductor fossa; **C**, interpretive drawing.

The maxilla is the largest bone of the upper jaw. It consists of a curved suborbital arm and a large postorbital expansion (Fig. 2). The dorsal margin of this expansion is incomplete. Roughly the upper one-third of the bone is missing with the exception of two small fragments immediately posterior to the jugal and suborbital. The ventral margin of the maxilla is ‘S’-shaped, and bears two series of teeth: large, widely spaced inner teeth and smaller, closely spaced outer teeth.

The dentary is the largest bone of the lower jaw (Figs 2B, 5A), covering almost its entire lateral surface. It bears at least 12 large conical teeth. The mandibular sensory canal extends along the ventral border of the dentary, and there is a foramen below the mentomeckelian that represents the exit of the trigeminal nerve to the inner jaw surface (Fig. 5). The ornamentation of this bone is similar to the other dermal bones of the skull, being only distinct due to the more robust ornamentation on the anterior portion of this bone. The angular and surangular represent the remaining bones on the external surface of the mandible. The angular is partially covered by the dentary but its ventral margin is visible along the posterior half of the dentary. The dentary partially overlaps the angular posteriorly, and tapers in this region. The surangular bone is poorly preserved, but represents a thin lamina dorsal to the angular and composes most of the external margin of the adductor fossa.

The primary jaw is only partially mineralized. The articular is ossified and rhomboidal, bearing two concave articular facets that mark the joint with the quadrate condyles (Fig. 5B). A small rugose ossification on the distalmost portion of the inner surface of the dentary might be a mentomeckelian ossification (Fig. 5C).

The internal wall of the lower jaw is presumably formed by the prearticular and the coronoids (Fig. 5B), although their boundaries cannot be discerned. The region interpreted as the prearticular consists of a vertical sheet of bone that forms the mesial wall of the adductor chamber and has a convex ventral margin. The anteriormost coronoids bear small conical teeth, but no teeth can be resolved more posteriorly in scans. The adductor fossa is large and triangular in dorsal view, bordered by the dentary anteriorly, the surangular and prearticular laterally and the articular posteriorly (Fig. 5C).

The palatoquadrate complex is partially preserved, and divisions between constituent ossifications are not apparent. Its shape broadly mirrors that of maxilla, with an expanded posterior blade and narrow subortbital ramus. At the junction of these two regions, the dorsal margin of the palatoquadrate bears a shallow embaument marking the position of the basipterygoid articulation. This is open dorsally, rather than being an enclosed fenestra (Fig. 5C). One series of teeth extends along the ventral margin of the palate. These teeth are intermediate in size between those of the two dentary tooth rows. It is unclear if this tooth row is restricted to the dermopalatines or are also borne by the ectopterygoid.

#### Operculo-gular Apparatus

The median gular is elongate and ellpisoidal, and bears a longitudinally oriented pit-line on the its external surface. The lateral gulars are similar in size to the median gular, but do not bear a pit-line. Both median and lateral gulars cover the anterior half of the intramandibular region (Fig. 6A-B). μCT reveals one extra pair of gulars buried within the matrix. Due to their position and shape they are herein described as extralateral gulars, laying behind the lateral gulars and extending until the first pair of branchiostegal rays (Fig. 6C-D).

**FIGURE 6.**
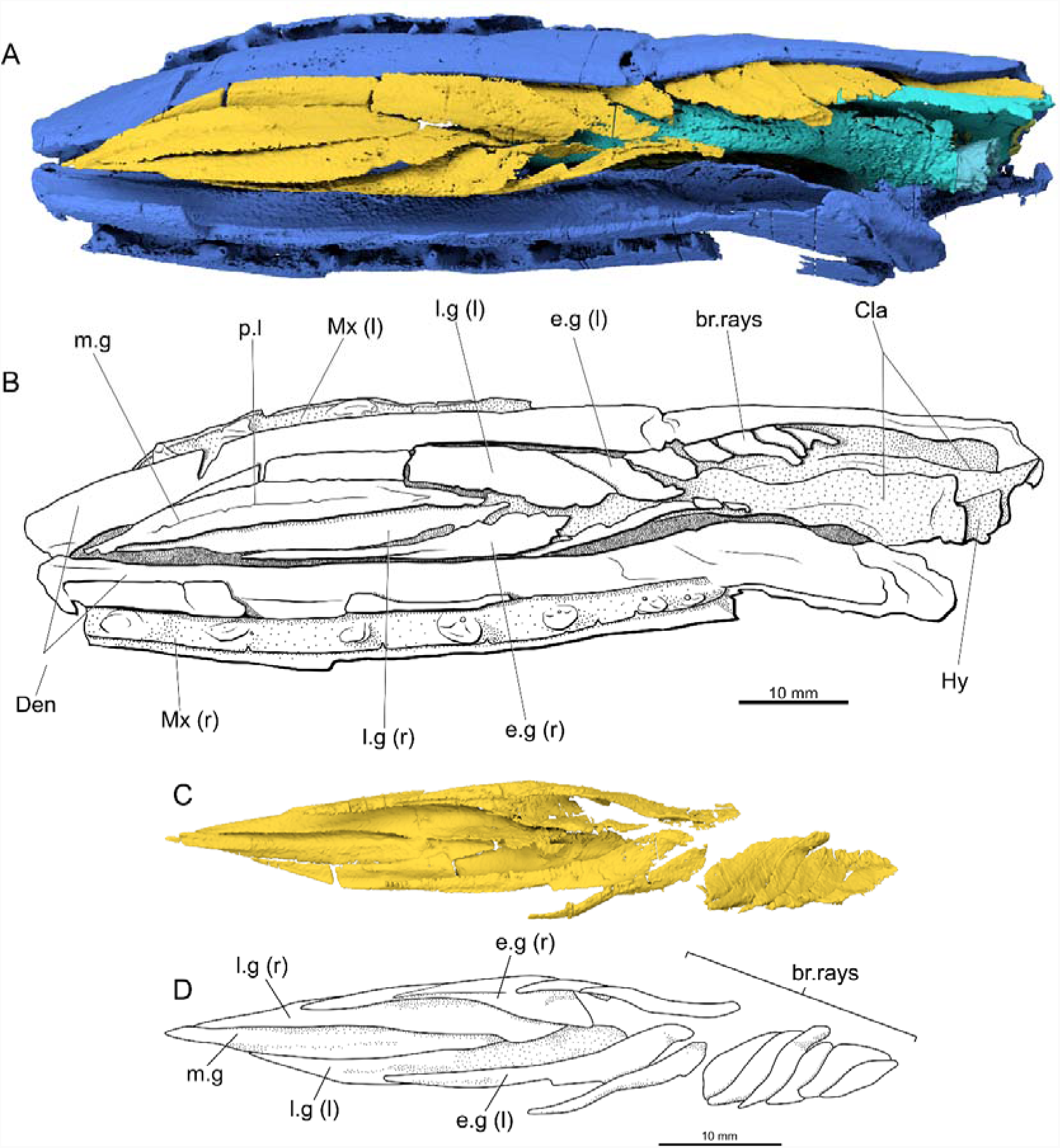
*Brazilichthys macrognathus*, DGM 1061-P, ventral view of jaws and intermandibular region. **A**, surface render based on μ-CT data; **B**, interpretive drawing of A; **C**, surface render of the gulars in dorsal view based on μ-CT data; **D**, interpretive drawing of C.

At least seven pairs of branchiostegal rays are partially preserved and visible externally. The posterior rays are partially broken, near the jaw articulation. The first pair is elongated but differs from the lateral and extralateral gulars by not being flattened. The exposed branchiostegal rays exhibit the same ornamentation pattern of other dermal bones of the skull: thin, wavy and closely arranged ridges.

The opercular series is almost completely missing, with the exception of the left opercle, which is preserved as a thin lamina behind the palatoquadrate complex (Fig. 3B). Due to the poor preservation of this element, it is impossible to provide accurate descriptions of the opercular series. Small fragments of a canal bearing overlapping the dorsal part of the palatoquadrate likely represent portions of the preoperculum, but are too incomplete to provide clear information on the bone.

#### Parasphenoid, Braincase and Associated Ossifications

The parasphenoid (Fig. 7A-F) coprises a long, slender anterior corpus that is expanded anterodorsally, likely for articulation with the vomer(s) and ethmoid region of the braincase, neither of which is well preserved. The anterior process of the parasphenoid is sub-triangular in cross-section. The ventral surface of the parasphenoid is smooth, with no evidence of a buccohypophysial foramen, any dentigerous area or ornamentation. However, individual denticles are likely beyond the resolution of the scan, so it is not possible to exclude the possibility that a denticle field was present. Stout dermal basipterygoid processes emerge from the lateral margin of the parasphenoid corpus immediately anterior to the broad ascending processes. Viewed dorsally, each basipterygoid process bears a small notch along its anterior margin at its junction with the body of the paraphenoid. The ascending processes expand dorsally, terminating with a straight margin. A shallow spiracular groove extends along the external surface of the left ascending process. There is no evidence of a posterior extension of the parasphenoid behind the ascending processes.

**FIGURE 7.**
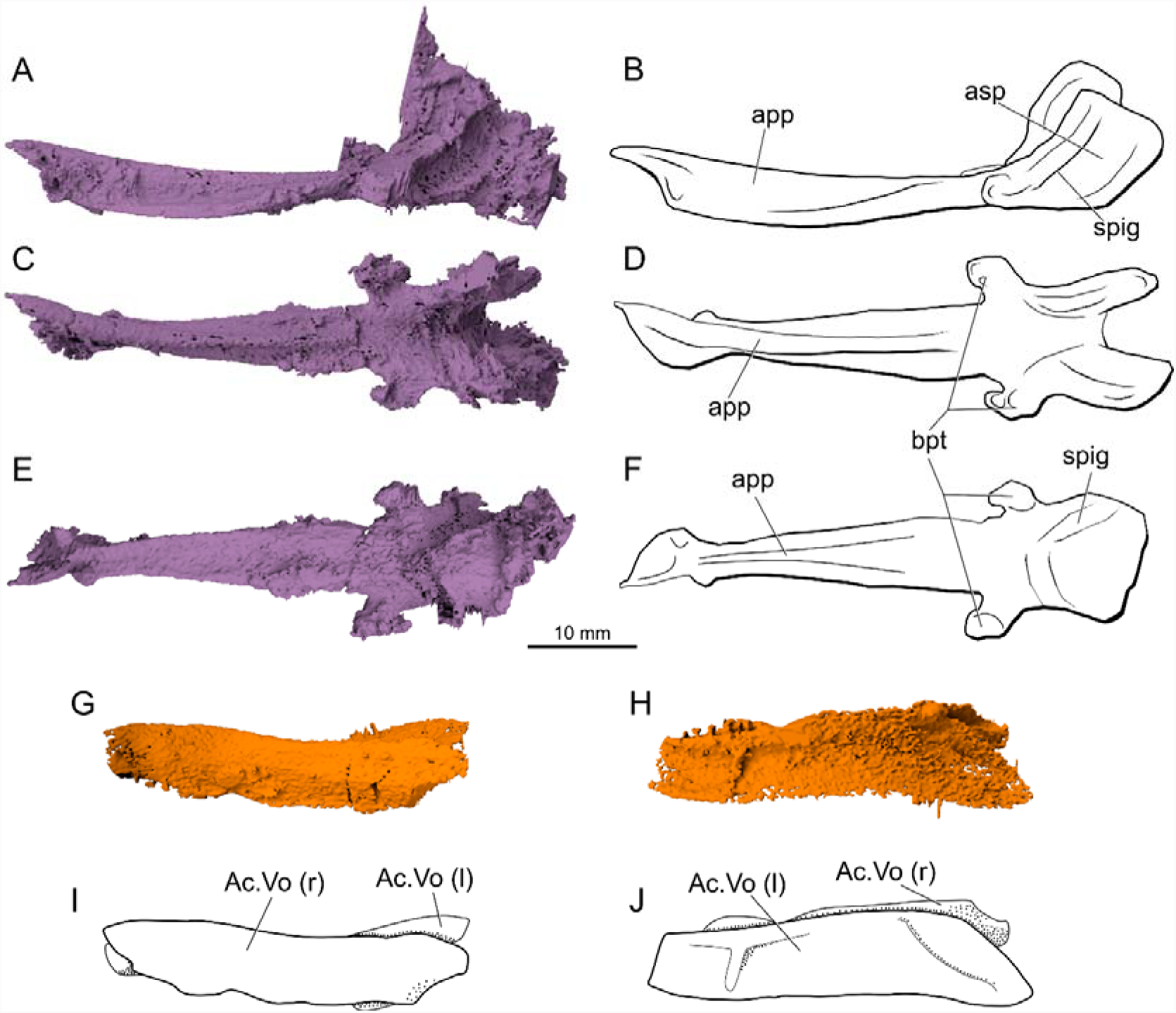
*Brazilichthys macrognathus*, DGM 1061-P, parasphenoid and accessory vomer. **A**, surface render in left-lateral view based on μ-CT data; **B**, interpretive drawing; **C**, surface render in dorsal view; **D**, interpretive drawing; **E**, surface render in ventral view; **F**, interpretative drawing; **G**, surface render in right-lateral view based on μ-CT data; **H**, interpretive drawing; **I**, surface render in left-lateral view; **J**, interpretive drawing.

Two long laminar bones preserved lateral to the parasphenoid represent accessory vomers (Fig. 7G-J). These plates contributed to the roof of the mouth in life, and would occupy the entire lateral margins of the anterior corpus of the parasphenoid. As the parasphenoid, these accessory vomers do not bear teeth large enough to be apparent in our scans. Compression of the skull resulted in a displacement of the right accessory vomer to the opposite side of the skull.

There is no trace of the neurocranium, and is assumed to be cartilaginous.

#### Hyoid and Branchial Arches

The left hyomandibula (Fig. 8A-D) is well preserved. It is strongly reclined, and the angle between its dorsal and ventral limbs is subtle. The dorsal limb is spatulate, with a compressed and proximal region representing the articular head. By contrast, the ventral limb is more cylindrical, with a rounded cross-section. Attached to the anterodorsal portion of the hyomandibula there is a fragment of what would be the preopercle (Fig. 8D), but the poor preservation of this element turns impossible its identification. The hyomandibular canal extends along the mesial surface of this element, exiting to the lateral surface by a foramen, before the expanded anterior surface of the hyomandibula. A sight dorsal expansion at the junction between the dorsal and ventral limbs of the hyomandibula represents a weakly developed opercular process. No dermohyal is preserved, and it was apparently not fused to the hyomandibula.

**FIGURE 8.**
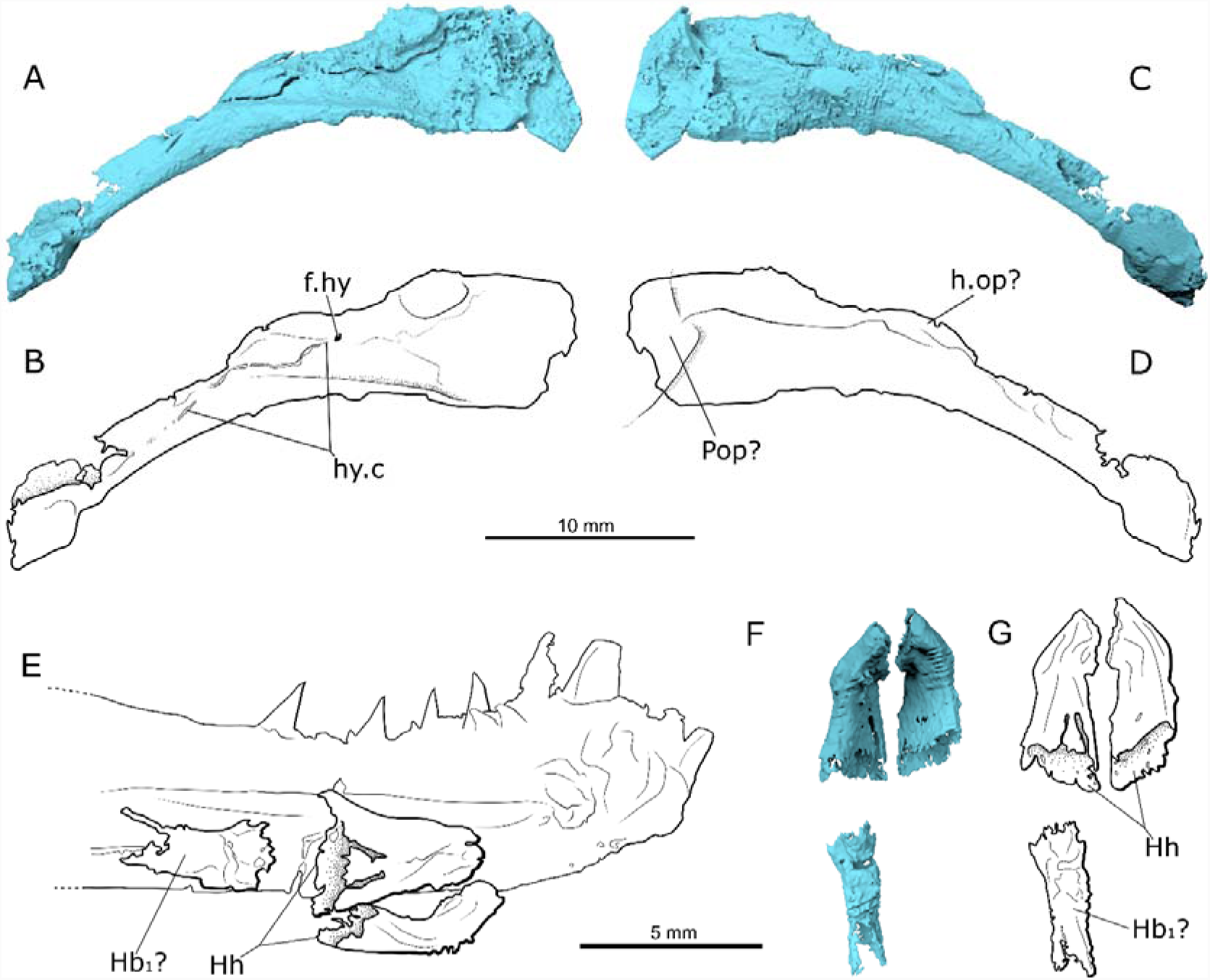
*Brazilichthys macrognathus*, DGM 1061-P, hyoid complex. **A**, surface render in mesial view based on μ-CT data; **B**, interpretive drawing; **C**, surface render in lateral view; **D**, interpretive drawing; **E**, hypohyals in right-lateral view shown in position relative to the mandible; **F**, render of ventral hyoid arch in dorsal view based on μ-CT data; **G**, interpretive drawing of F.

The hypohyals (Fig. 8E-G) are well preserved, located in the distal portion of the ventral surface of the lower jaw. They are cylindrical and expand posteriorly, and curve toward one another along the midline. There is no evidence of a mineralized basibranchial.

Preserved components of the branchial skeleton (Fig. 9) are located in the posterior half of the skull and consist of four pairs of long rods that probably represent ceratobranchials. There are two short epibranchials that do not show an evidence of developed uncinate processes. The hypobranchials are long, expanded anteriorly to for their articulation with the unpreserved basibranchial. Other smaller elements are present, but fragmentation and displacement makes identifications difficult.

**FIGURE 9.**
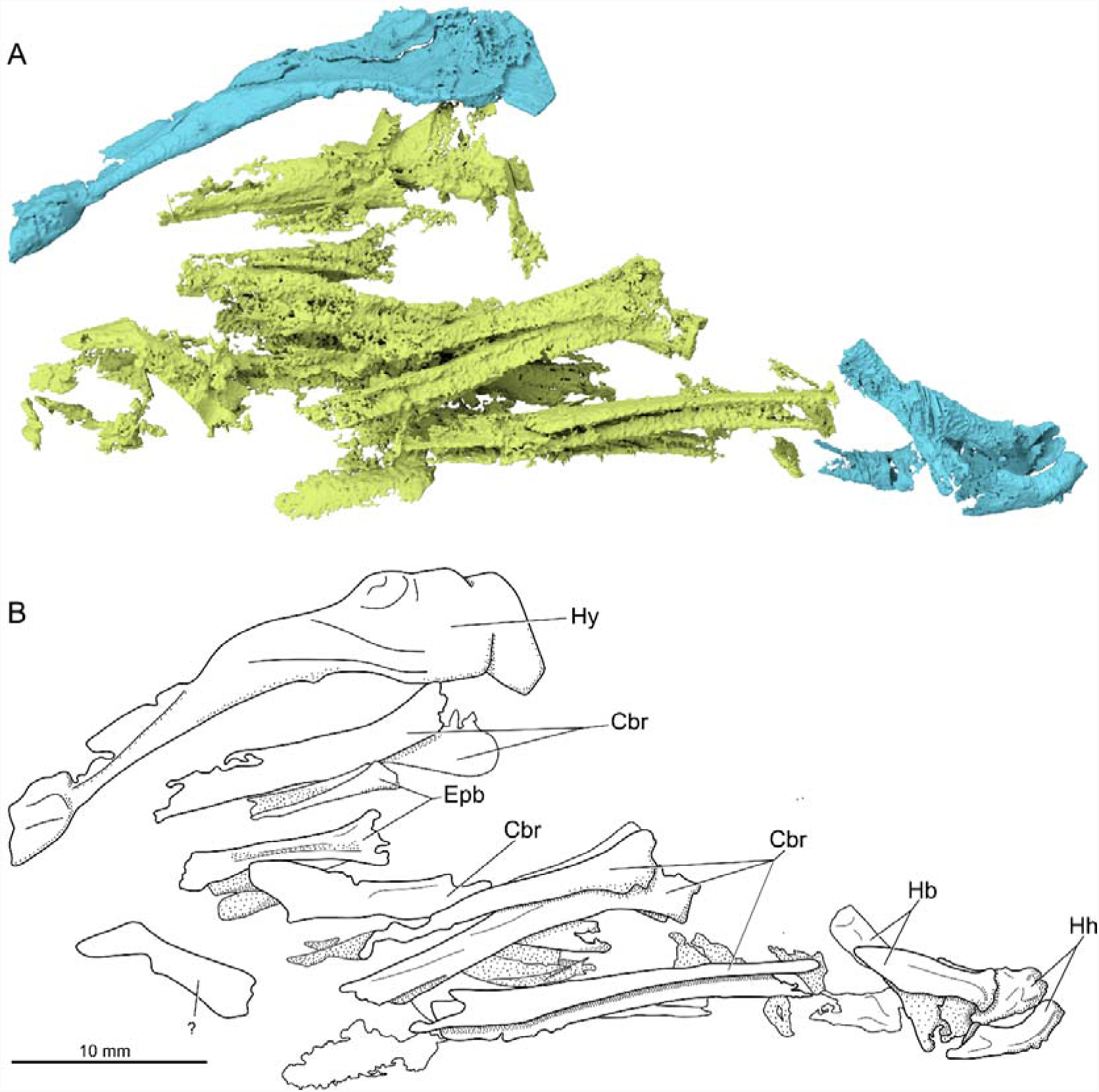
*Brazilichthys macrognathus*, DGM 1061-P, hyoid and branchial arches in right-lateral view. **A**, surface render based on μ-CT data; **B**, interpretive drawing.

#### Shoulder Girdle

The clavicles (Fig. 10A-B) bound the branchial skeleton ventrally. Each clavicle has an elongated anterior ramus and terminates in a lamina with a triangular profile in dorsal view. There is a displaced, ellipsoidal bone between the branchial rays and dorsal to the clavicles that could be a poorly preserved interclavicle (Fig. 10C), however it is not possible to accurately identify this bone, and therefore it was not coded as present in the phylogenetic matrix. Other components of the shoulder girdle are not preserved. However, one small rhomboid element located posteriorly to the end of the clavicles could be interpreted as the first fin rays that fused to form this rigid structure.

**FIGURE 10.**
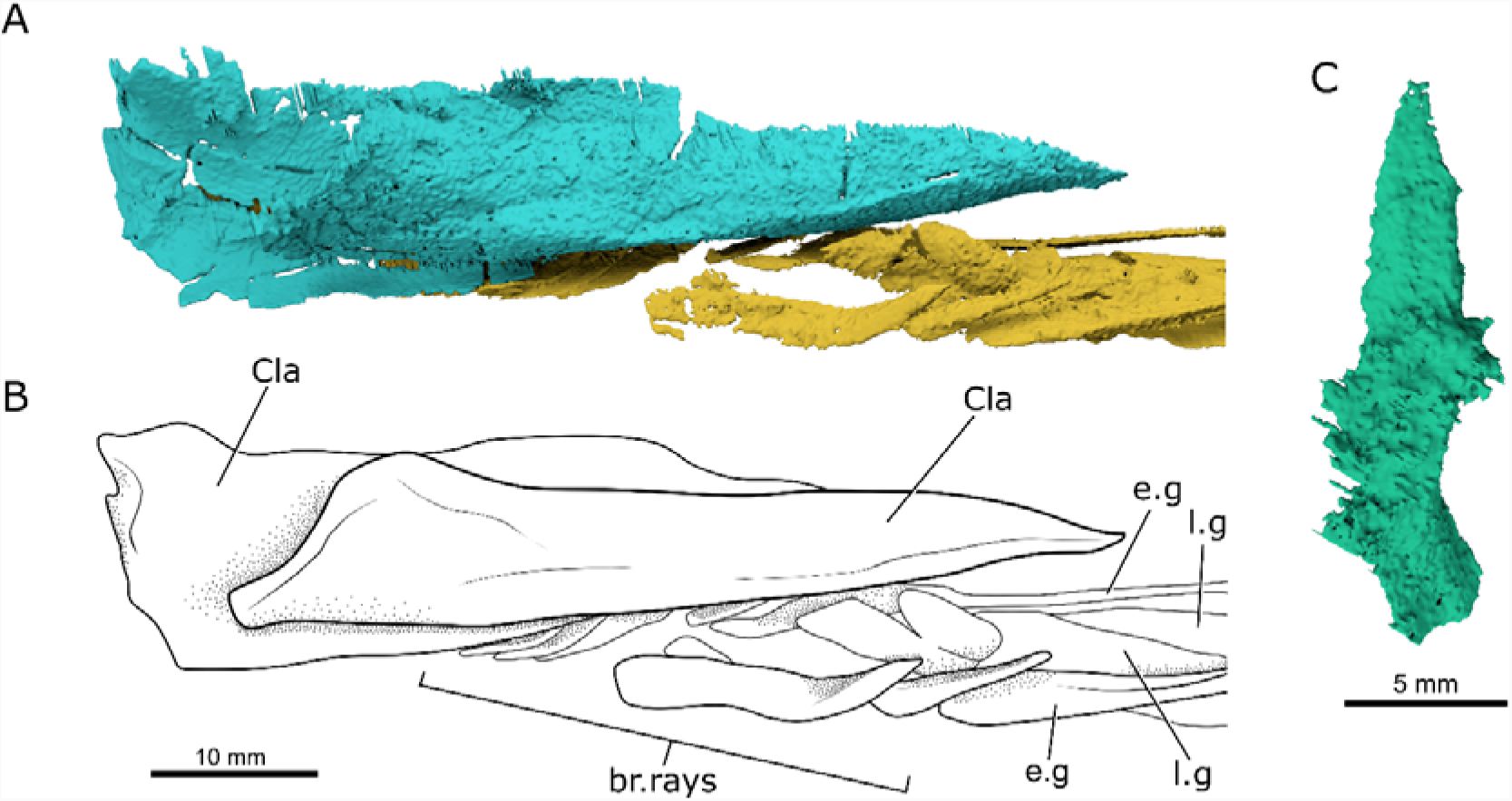
*Brazilichthys macrognathus*, DGM 1061-P, clavicles in ventral view. **A**, surface render based on μ-CT data; **B**, interpretive drawing. **C**, surface render of possible interclavicle.

### Phylogenetic results

#### Parsimony Analysis

The parsimony analysis including *Brazilichthys* recovered four equally parsimonious trees (length = 1325 steps; consistency index: 0.22; retention index: 0.64). The strict consensus is well resolved (Fig. 11), showing conventional actinopterygian, sarcopterygian and chondrichthyan clades. Devonian actinopterygians form a grade on the actinopterygian stem. *Meemannia*, a clade uniting the species of *Cheirolepis*, and a clade uniting *Osorioichthys* and *Tegeolepis* are earliest diverging actinopterygian lineages, with Middle-Late Devonian taxa (*Donnrosenia, Howqualepis, Mimipiscis, Gogosardina, Raynerius, Moythomasia*) forming a more crownward clade (Bremer decay index [BDI] = 2). *Brazilichthys* is resolved crownward of all Devonian taxa, in a polytomy with two other clades (BDI = 2). The first of these includes most late Paleozoic taxa conforming to a generalized ‘palaeoniscoid’ habitus and is poorly supported (BDI = 1). The second unites *Saurichthys* and *Australosomus* with the actinopterygian crown. *Discoserra, Ebenaqua, Platysomus, Amphicentrum, Styracopterus* and *Fouldenia* are the most highly nested Paleozoic taxa, and are placed on the chondrostean stem (BDI = 3). Among extant actinopterygians, chondrosteans and cladistians represent successively more remote outgroups to neopterygians.

**FIGURE 11.**
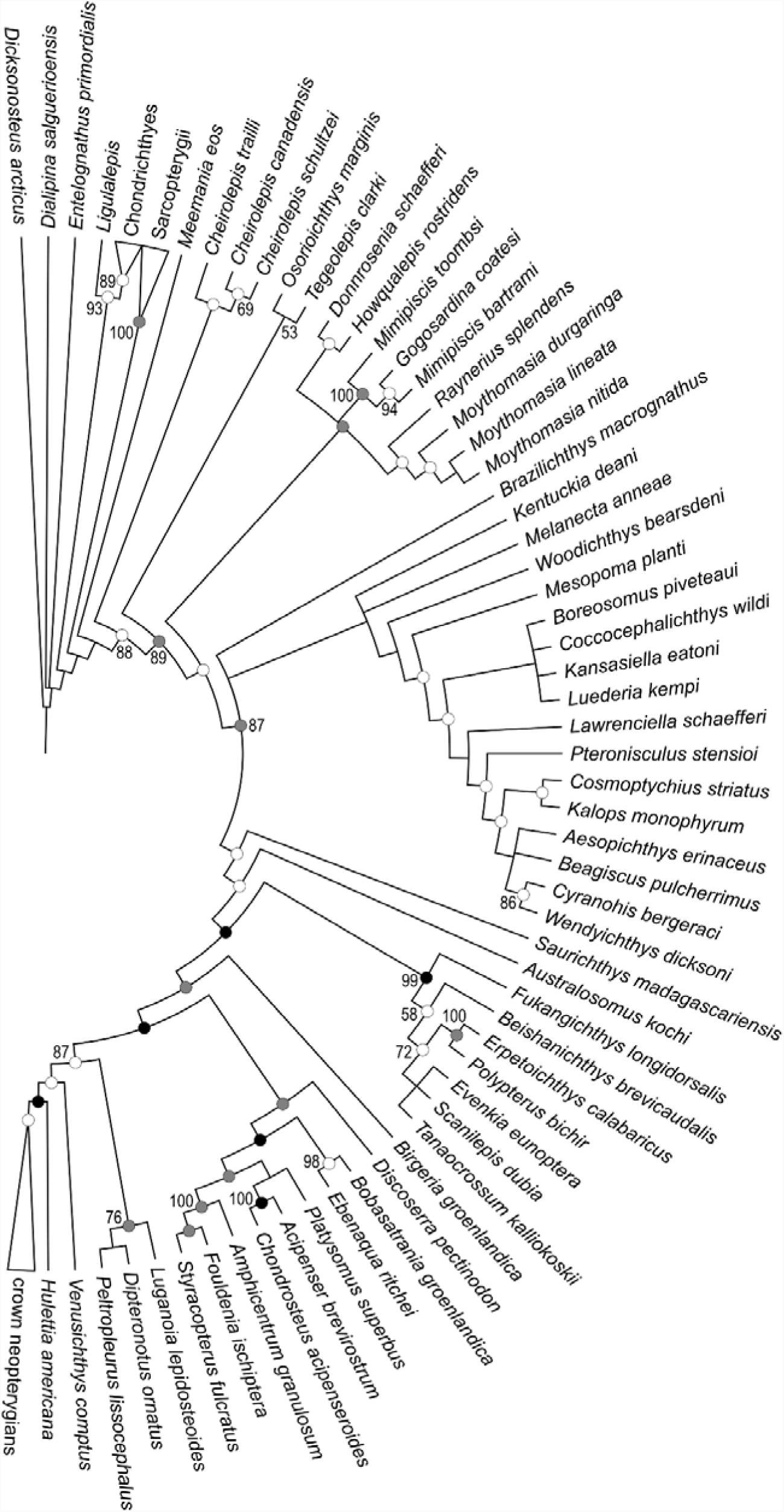
Phylogenetic hypothesis of early actinopterygian interrelationships, based on maximum parsimony analysis of 265 morphological characters (from Giles et al., 2017). Tree length: 1326 steps; consistency index: 0.22; retention index: 0.64. Node colors represent Bremer support values: white = 1, light gray = 2, black ≥ 3. Numbers represent posterior probabilities (x 100), and are only shown for nodes found in the majority-rule consensus of the posterior distribution of trees in the Bayesian analysis.

The following characters are optimized as arising along the branch subtending the clade including *Brazilichthys* plus all other post-Carboniferous taxa: dermopterotic carying lateral line canal between dermosphenotic and skull roof (36; 0 – 1); absence of a distinct posterior ramus of the dermosphenotic (56; 1 – 0); presence of two coronoids (91; 1 – 3); presence of median posterior myodome (136; 0 – 2); spiracle enclosed in canal (138; 1 – 2); presence of fossa bridgei (143; 0 – 1); absence of anterolaterally divergent olfactory tracts (180; 1 – 0); optic lobes of same width or wider than cerebellum (184; 0 – 1); optic tectum divided into bilateral halves (186; 0 – 1); presence of cerebellar corpus (187; 0 – 1); 193 (crus commune ventral to endocranial roof (193; 0 – 1); presence of opercular process of hyomandibula (211; 0 – 1); and absence of interclavicle (233; 0 – 1). These synapomorphies were found using both parsimony and likelihood ancestral state reconstructions. It is interesting to notice that despite being recovered as a synapomorphy, the presence of two coronoids has a low probability (0.47). However, of these synapomorphies only the absence of a distinct posterior process of the dermosphenotic and presence of an opercular process of the hyomandibula can be shown for *Brazilichthys*, and both characters show some degree of homoplasy. Sarcopterygians have a dermosphenotic (sarcopterygian intertemporal; Schultze, 2008) that lack a posterior ramus, while Devonian actinopterygians bear a distinct posterior ramus of the bone. This posterior extension is lost in the clade uniting *Brazilichthys* and all post-Devonian actinopterygians, with multiple reversals within the group (e.g. *Beagiascus, Wendyichthys, Cyranorhis, Birgeria, Bobasatrania* and some neopterygians). With respect to the hyomandibula, sarcopterygians, chondrichthyans and the earliest actinopterygians show absence of an opercular process. The node including *Brazilichthys* and all other post-Devonian is united by presence of opercular process, with reversals in *Acipenser, Amphicentrum, Birgeria, Chondrosteus*, and *Kalops*.

#### Bayesian Analysis

The Bayesian analysis shows a similar result to the parsimony analysis presented above. Major differences, described in more detail below, include: the degree of phylogenetic resolution among Paleozoic actinopterygians; the content of the chondrostean stem; and the interrelationships of neopterygians. With the exception of those subtending (1) Cladistia inclusive of scanilepiforms and (2) *Condrosteus* plus *Acipenser*, nodes that are well-supported in the parsimony analysis parsimony analysis (BDI ≥ 3) are not represented in the Bayesian tree.

As in the parsimony tree, *Meemannia* and *Cheirolepis* represent the earliest-diverging ray-finned fishes, although the monophyly of the latter genus is not supported (Fig. 11). All other Devonian actinopterygians are placed crownward of *Cheirolepis*, although their relationships are unresolved apart from a poorly supported clade uniting *Osorioichthys* and *Tegeolepis* (Bayesian posterior probability [BPP] = 0.55) and a better supported clade containing *Mimipiscis* with *Gogosardina* nested within that genus (BPP = 1.00). Post-Devonian actinopterygians, including *Brazilichthys*, form a clade (BPP = 0.87). The relationships among most Paleozoic members of this group are not resolved. *Brazilichthys* is placed in a polytomy with the actinopterygian crown (BPP = 0.52), eurynotiforms (BPP = 1.00), a clade uniting *Wendyichthys* and *Cyranorhis* (BPP = 0.86), and over a dozen species-level lineages of unresolved affinity. Within the actinopterygian crown, the relationships between the cladistian, chondrostean, and neopterygian lineages are unresolved. Scanilepiforms are placed as stem cladistians with high support (BPP = 0.99), but *Birgeria* is the only member placed on the chondrostean stem (BPP = 0.80) apart from the phylogenetically uncontroversial *Chondrosteus*. *Platysomus* and a clade uniting *Discoserra, Ebenaqua*, and *Bobasatrania* (BPP = 83) are placed as the deepest diverging lineages on the neopterygian stem, and include the only Paleozoic members of the actinopterygian crown.

The placement of *Brazilichthys* crownward of all Devonian actinopterygians (Fig. 11) is supported by the same characters as in the parsimony solution. Ancestral state reconstruction using maximum likelihood and parsimony provides a similar picture of character support, the exception that the presence of three coronoids (probability = 0.48)--rather than two (probability = 0.28)--representing the most probable state at the base of this post-Devonian radiation. This uncertainty over this character was also present in the ancestral state reconstruction over the parsimony topology, as mentioned above. Ancestral-state reconstruction also suggests additional synapomorphies of the post-Devonian clade not indicated by parsimony mapping. Of these, characters 141 (presence of dermal component of basipterygoid process (141; 0 - 1) and absence of a multifid anterior margin of the parasphenoid (173; 1 - 0) and absence of buccohypophyseal canal piercing parasphenoid (174; 0 - 1) are the only ones that can be confirmed for *Brazilichthys*. A complete list of synapomorphies for the node uniting *Brazilichthys* and post-Devonian taxa is available in Supplementary Data 4.

## DISCUSSION

### The Interrelationships of Early Actinopterygians: How Far from Consensus?

Until relatively recently, the dominant systematic paradigm for early actinopterygian systematics placed nearly all Paleozoic taxa within the crown. This arrangement began with relatively abbreviated, hand-constructed solutions presented by Patterson (1982) and Gardiner (1984), followed by more extensive (Gardiner and Schaeffer, 1989) and algorithmic analyses (Coates, 1999; Gardiner et al., 2005; Xu et al., 2014). However, several cladistic analyses have suggested that many Paleozoic forms are stem actinopterygians (Cloutier and Arratia, 2004), a viewpoint recently reinforced by the confirmation that scanilepiforms represent stem cladistians (Giles et al., 2017). The latter discovery reveals that many of the apparently primitive features of extant polypterids are secondary, with the consequence that many fossils previously placed with the actinopterygian crown have shifted to the stem. Despite the addition of new taxa, modification of specific aspects of coding, or the investigation of new characters, this pattern has emerged consistently in re-analysis of datasets that trace their origin to that of Giles *et al.* (2017; e.g. Argyriou et al., 2018; Latimer and Giles, 2018). Apart from the placement of most Paleozoic taxa on the actinopterygian stem, this set of results is characterized by a series of similar features. Apart from uncontroversial aspects of relationships within the crown (e.g. the monophyly of major living lineages) or long-established patterns on the stem (e.g. the early divergence of *Cheirolepis*) these include the monophyly of all post-Devonian taxa and the placement of one or more deep-bodied Paleozoic lineages within the actinopterygian crown. Both patterns merit further discussion.

With respect to the monophyly of post-Devonian taxa, this ‘phylogenetic bottleneck’ could be interpreted as the radiation of a single surviving lineage after the end-Devonian extinction, mirroring models of diversification sometimes proposed for birds (Prum et al. 2016) and mammals (O’Leary et al., 2013) in the early Cenozoic. While there is undoubtedly ample evidence for taxonomic (Gardiner, 1993), morphological (Sallan and Friedman, 2012), and ecological (Friedman et al. 2019) diversification of actinopterygians in the Carboniferous relative to the Devonian, the recent interpretation of the earliest Carboniferous *Arvonichthys* as the nested within the clade uniting the Middle-Late Devonian *Howqualepis, Donnrosenia, Moythomasia*, and *Mimipiscis* complicates this model of diversification from a single surviving lineage (Wilson et al., 2018). Further study of Famennian and Tournaisian study is vital for clarifying patterns of actinopterygian turnover at the end-Devonian extinction.

With respect to the placement of deep-bodied lineages of Paleozoic actinopterygians within the crown, the pattern is more complicated than this general summary suggests. The exact placement of these lineages varies between studies, as well as within studies as a function of the approach to phylogenetic inference (i.e. parsimony versus Bayesian). Only *Discoserra, Platysomus*, and *Ebenaqua* (along with the similar Triassic *Bobastrania*) are always placed within the crown, with eurynotiforms vacillating between the crown and stem. When any taxa are placed within the crown, they lie either on the chondrostean (e.g. Wilson et al., 2018) or neopterygian (e.g. Argyriou et al., 2018) stem. More broadly, this reflects a common pattern across this set of hypotheses of early actinopterygian interrelationships: overall similarities at very coarse scales, but substantial disagreement at lower levels. The fact that these relationships vary with relatively minor modifications to either the dataset of mode of phylogenetic inference suggests that caution should be applied in interpreting the significance of branching patterns among Paleozoic actinopterygians at medium to fine phylogenetic scales. It is worth reiterating that deficient anatomical knowledge of most Permo-Carboniferous fishes appears to be a major contributor to their phylogenetic instability (Giles et al., 2017). We therefore remain optimistic that more detailed studies of internal structure in fossils of this age (Coates and Tietjen, 2019; Friedman et al. 2019), including our own redescription of *Brazilichthys*, will prove helpful in solidifying our understanding of relationships among Paleozoic actinopterygians. This will represent an important phylogenetic foundation for the increasing number of studies addressing macroevolutionary patterns in Paleozoic fishes, many of which have been conducted in a non-phylogenetic framework (e.g. Anderson et al., 2011; Sallan and Giamberti, 2015).

### The Relationships of *Brazilichthys* and Ecomorphologically Similar Taxa

#### Assessment of previous hypotheses

Prior to this study, *Brazilichthys* had not been included in a formal phylogenetic analysis. Instead, interpretations of its systematic position have been based on perceived similarities with other early ray-finned fishes. In their original description of *Brazilichthys*, Cox and Hutchinson (1991) suggested a relationship with Acrolepidae, based on a suggestion from B. G. Gardiner citing the “form of the teeth and the elongated maxilla.” As originally delimited by Aldinger (1937), acrolepids contained a diverse set of generalized early actinopterygians ranging in age from Devonian to Jurassic in age (*Watsonichthys, Acrolepis, Acropholis, Plegmolepis, Reticulolepis, Hyllingea, Boreosomus, Acrorhabdus, Diaphorognathus, Pytcholepis*, and *Stegotrachelus*). There is little in his diagnosis that appears unique to the group, or indeed which is shared among all its members, with strongly contrasting states being described as diagnostic for the family (e.g. either few or many branchiostegal rays, either a large median gular or a small median gular, either a vertical suspensorium or an oblique suspensorium, and so on). Gardiner (1963) referred to Aldinger’s diagnosis with no modifications, and Stamberg (1991) only provided modest amendments. We have no confidence that Acrolepidae, as construed by these authors, represents a monophyletic group. Here we restrict our comparisons with *Acrolepis*, the type acrolepid. Numerous species have been questionably referred to *Acrolepis* (see summary in Aldinger, 1937), with many removed to other genera (e.g. ‘*Acrolepis*’ *laetus*, which is now recognized as a species of *Pteronisculus*; Schaeffer and Mangus, 1976). We therefore further restrict our comparisons to the type species, *A. sedgwicki* from the Permian Marl Slate of England. Descriptions of this taxon are sparse, with published interpretive drawings secondary representations derived from unpublished sources (e.g. Aldinger, 1937: fig. 74, from T. S. Westoll’s unpublished 1934 dissertation from the University of Durham on the Permian ‘paleoniscoids’ of the Marl Slate; Gardiner and Schaeffer 1989: fig. 9, attributed only to an unpublished drawing by P. Hutchinson). Apart from shared generalities, there appears to be little evidence that *Acrolepis* and *Brazilichthys* might be closely related. Derived features shared between these taxa (e.g. absence of a posterior process of the dermosphenotic, the presence of at least one suborbital) are widely distributed among post-Devonian actinopterygians. Unfortunately, *Acrolepis* has not been included in a formal phylogenetic analysis and is too incomplete to include here. While we regard *Acrolepis* and *Brazilichthys* as plausibly branching from the same general region of actinopterygian phylogeny—on the actinopterygian stem crownward of Devonian forms—we see no particular reason to regard them as especially closely related to one another to the exclusion of other Permo-Carboniferous paleopterygians. Indeed, there are several broad disagreements between known skeletal anatomy between *Acrolepis* and *Brazilichthys*, mostly related to the geometry of the suspensorium and the proportions of the maxilla and mandible. We would not be surprised if more detailed understanding of internal anatomy of *Acrolepis* highlighted further differences from what we report here in *Brazilichthys*.

More recently, Romano and Brinkmann (2009) suggested a possible relationship between *Brazilichthys* and *Birgeria*. Like the interpretation proposed by Cox and Hutchinson (1991) these authors made this proposal on the basis of personal communication with a colleague (R. J. Mutter) with no reference to specific anatomical evidence. However, it seems likely that this comparison derives from the long mandible and cheek region and the posteriorly directed suspensorium apparent in both taxa, combined with a mistaken interpretation of *Brazilichthys* as being late—rather than mid—Permian in age and thus a more immediate stratigraphic antecedent of *Birgeria*. However, apart from such generalities there are no obvious synapomorphies that would unite *Birgeria* and *Brazilichthys* to the exclusion of a variety of other early actinopterygians. Indeed, our new findings about the internal anatomy of *Brazilichthys* reveal several pronounced contrasts with *Birgeria*. Among the most conspicuous relate to the dermal basipterygoid process of the parasphenoid (present in *Brazilichthys*, but absent in *Birgeria*; Nielsen, 1949: fig. 70), the posterior stalk of the parasphenoid (absent in *Brazilichthys*, but present in *Birgeria*; Nielsen, 1949: fig. 70), the posterior margin of the orbit (defined by a single jugal ossification in *Brazilichthys*, but comprising a chain of ossicles in *Birgeria*; Nielsen, 1949: fig. 69), and the geometry of the hyomandibula (subequal dorsal and ventral limbs in *Brazilichthys*, but a much longer dorsal limb in *Birgeria*; Nielsen, 1949: fig. 72). Our analyses strongly reject the hypothesis of a close relationship between *Birgeria* and *Brazilichthys*, placing these genera far apart from one another. Regardless of the mode of phylogenetic inference, *Brazilichthys* is always resolved as a stem actinopterygian crownward of all Devonian taxa, while *Birgeria* lies within the crown radiation as either a stem chondrostean (Bayesian) or stem actinopteran (parsimony).

#### The Problem of Large, Predatory Paleopterygians

Past attempts to determine the likely phylogenetic placement of *Brazilichthys* reflect a broader problem relating to the systematics of early actinopterygians more generally. While some early actinopterygians form reasonably well circumscribed groups supported by clear synapomorphies (e.g. eurynotiforms, platysomids, haplolepids), the vast majority of early ray-finned fishes present an outwardly similar suite of characters of the dermal skeleton with only minor variations between them. Within this assemblage of anatomically generalized taxa, there are a variety of forms characterized by reasonably large size (cranial length ca. 50 mm or larger) and the presence of a proportionately large row of inner dentary teeth, traits that suggest a predatory—and likely piscivorous—ecology, and inference supported in some cases by the presence of gut contents (e.g. Gardiner 1963). Permo-Carboniferous taxa of this broad ecological gestalt are divided between a series of notional families in addition to Acrolepidae, including Pygopteridae, Cosmoptychiidae, and Rhabdolepidae. Excluding the monogeneric Rhabdolepidae, all of these groups—like Acrolepidae—are of questionable monophyly. The association of *Brazilichthys* with both *Birgeria* and acrolepids reflects a series of general (oblique suspensorium) and more specialized (proportionally large teeth) features that equally characterize these other Permo-Carboniferous taxa. However, it is not possible to discern the mutual relationships of these ecomorphologically similar Paleozoic fishes in light of data currently available, which is almost entirely restricted to cursory descriptions of the external dermal skeleton. However, many of these taxa are known from relatively uncrushed, three-dimensionally preserved skulls (*Cosmoptychius* and *Nematoptychius* from Wardie, Scotland; *Rhabdolepis* from Lebach, Germany). We are hopeful that additional details of the internal anatomy of some of these taxa will help to establish their relative phylogenetic relationships, providing some insight into the number of times this particular ecomorph arose in early actinoperygian history. This, in turn, is relevant for understanding patterns of actinopterygian trophic diversification in the Carboniferous (Sallan and Friedman 2012). Such an effort would also help to reduce the seemingly overwhelming early actinopterygians by identifying smaller candidate clades that could reduce the overall scale of the phylogenetic problem through stepwise approach to the resolution of relationships (cf. eurynotiforms; Sallan and Coates 2013; Friedman et al. 2019).

## CONCLUSIONS

Re-examination of the holotype of the mid-Permian (Artinskian-Kungurian) actinopterygian *Brazilichthys macrognathus* from Brazil reveals considerable new anatomical information. Revisions of details of external anatomy relative to the initial description by Cox and Hutchinson include reinterpretation of a supposed sclerotic as a dermosphenotic and identification of three rather than two bones composing the infraorbital series. A median rostral element previously described appears to be missing, possibly due to damage to the specimen. μ-CT provides important new information on internal anatomy not accessible to earlier researchers. This reveals the palate (including accessory vomers), the inner surface of the mandible, the parasphenoid, the hyoid arch, disrupted branchial arches, and portions of the clavicles. The braincase is unmineralized. Overall, the cranial anatomy of *Brazilichthys* shows generalized conditions for actinopterygians, but the presence of derived features like the presence of an ascending process and dermal basipterygoid processes of the parasphenoid, a suborbital, an opercular process of the hyomandibula, and absence of a long posterior process of the dermosphenotic suggest that it branches crownward of the very earliest diverging ray-finned fishes. Formal phylogenetic analyses place *Brazilichthys* crownward of all Devonian actinopterygians. The parsimony solution resolves *Brazilichthys* as the sister lineage to all remaining members of the post-Devonian actinopterygian clade, while the Bayesian analysis places the genus in a large polytomy at the base of this radiation. We reject previous hypotheses for the placement of *Brazilichthys*, which are based largely on overall similarities reflecting primitive actinopterygian conditions. This genus is retained as the only member of Brazilichthyidae pending more detailed analysis of large-bodied Permo-Carboniferous fishes with similarly generalized external anatomy and jaws suggestive of a predatory ecology.

## ACKNOWLEDGMENTS

The authors are thankful to the Centro de Pesquisa de Recursos Minerais – CPRM, Departamento Nacional de Produção Mineral – DNPM and R.M. da Silva for allowing the loan of DGM 1061-P. We thank the Laboratório de Instrumentação Nuclear, R.T. Lopes and A. Machado for conducting the CT-scan of the analyzed specimen. Acknowledgements are needed for L.M. Diele-Viegas for kindly reviewing an early version of the manuscript and S. Giles for assistance on the 3D rendering and taxonomy. RF also acknowledges Coordenação de Aperfeiçoamento de Pessoal de Nível Superior – CAPES and Fundação de Amparo à Pesquisa do Rio de Janeiro – FAPERJ for the scholarships that funded this research. VG is supported by Conselho Nacional de Desenvolvimento Científico e Tecnológico - CNPq and have an UERJ/FAPERJ Prociência grant.

